# The dynamic chromatin architecture of the regenerating liver

**DOI:** 10.1101/664862

**Authors:** Amber W. Wang, Yue J. Wang, Adam M. Zahm, Ashleigh R. Morgan, Kirk J. Wangensteen, Klaus H. Kaestner

## Abstract

**Background & Aims:** The adult liver is the main detoxification organ and is routinely exposed to environmental insults but retains the ability to restore its mass and function upon tissue damage. However, massive injury can lead to liver failure, and chronic injury causes fibrosis, cirrhosis, and hepatocellular carcinoma. Currently, the transcriptional regulation of organ repair in the adult liver is incompletely understood.

**Methods:** We isolated nuclei from quiescent as well as repopulating hepatocytes in a mouse model of hereditary tyrosinemia, which recapitulates the injury and repopulation seen in toxic liver injury in humans. We then performed the ‘assay for transposase accessible chromatin with high-throughput sequencing’ (ATAC-seq) specifically in repopulating hepatocytes to identify differentially accessible chromatin regions and nucleosome positioning. Additionally, we employed motif analysis to predict differential transcription factor occupancy and validated the *in silico* results with chromatin immunoprecipitation followed by sequencing (ChIP-seq) for hepatocyte nuclear factor 4α (HNF4α) and CCCTC-binding factor (CTCF).

**Results:** Chromatin accessibility in repopulating hepatocytes was increased in the regulatory regions of genes promoting proliferation and decreased in the regulatory regions of genes involved in metabolism. The epigenetic changes at promoters and liver enhancers correspond with regulation of gene expression, with enhancers of many liver function genes displaying a less accessible state during the regenerative process. Moreover, increased CTCF occupancy at promoters and decreased HNF4α binding at enhancers implicate these factors as key drivers of the transcriptomic changes in replicating hepatocytes that enable liver repopulation.

**Conclusions:** Our analysis of hepatocyte-specific epigenomic changes during liver repopulation identified CTCF and HNF4α as key regulators of hepatocyte proliferation and regulation of metabolic programs. Thus, liver repopulation in the setting of toxic injury makes use of both general transcription factors (CTCF) for promoter activation, and reduced binding by a hepatocyte-enriched factor (HNF4α) to temporarily limit enhancer activity.

**Graphical Abstract:** 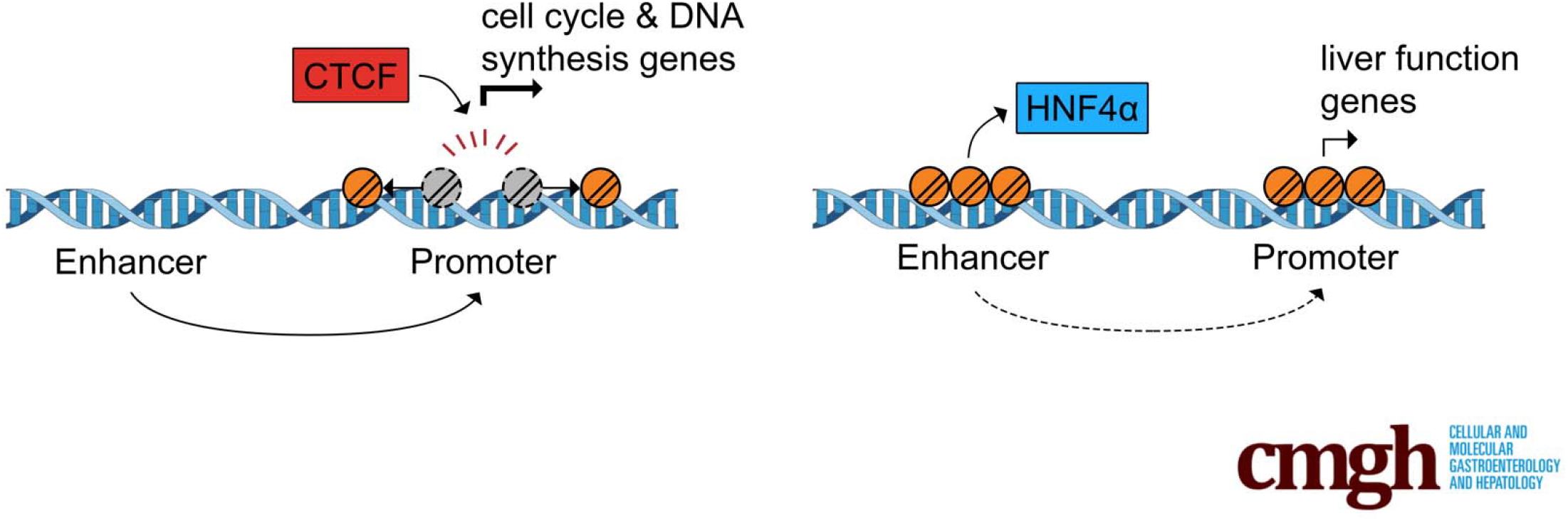

## INTRODUCTION

As the central metabolic organ in vertebrates, the liver regulates carbohydrate, protein, and lipid homeostasis, metabolizes nutrients, wastes, and xenobiotics, and synthesizes bile, amino acids, coagulation factors, and serum proteins^1^. To prevent acute liver failure upon exposure of harmful toxins, the liver has maintained an extraordinary ability to effectively restore its mass and function, in which the normally quiescent mature hepatocytes rapidly re-enter the cell cycle and divide^2^. Nonetheless, failure of regeneration can occur after exposure to harmful metabolites and environmental toxins, as often seen with the overconsumption of acetaminophen and alcohol^3^. Hence, understanding the genetic networks regulating the regenerative process can have an immense impact on the development of novel therapeutic strategies to treat acute liver failure.

The *Fah* null mouse model of human hereditary tyrosinemia type I provides a unique system to study the hepatocyte replication process after acute liver injury. These mice lack the fumarylacetoacetate hydrolase (FAH) enzyme, which is essential for normal tyrosine catabolism, and results in the accumulation of toxic intermediates followed by hepatocyte cell death^4,5^. *Fah*^−/−^ mice can be maintained in a healthy state by supplementation with the drug 2-(2-nitro-4-trifluoromethylbenzoyl)-1,3-cyclohexanedione (NTBC) in the drinking water which inhibits an upstream enzymatic step that prevents toxin production^4^. Alternatively, gene therapy that utilizes hydrodynamic tail-vein injection and the *Sleeping Beauty* transposon system to restore *Fah* expression can rescue these mice^6,7^. When a small fraction (0.1-1%) of hepatocytes come to express FAH following removal of NTBC, these hepatocytes competitively repopulate the liver in the context of injury through clonal expansion. Furthermore, this method allows lineage-tracing of repopulating hepatocytes since only those with stable FAH expression can expand and repopulate the injured parenchyma^7,8^.

Eukaryotic DNA is highly organized and structured into compact chromatin to allow tight transcriptional control. Transcriptional regulation can be broadly categorized into two integrated layers: (1) transcription factors and the transcriptional machinery, and (2) chromatin structure and its regulatory proteins^9^. Expression of genes targeted by transcription factors depends on the binding affinity to specific target DNA recognition sequences, combinatorial assembly with other cofactors, concentration of the factor, and post-translational modifications that affect protein localization^10^. The chromatin landscape is governed by DNA methylation, nucleosome properties, histone modifications, and intra- and interchromosomal interactions^10^. Establishing the relationship of chromatin structure, transcriptional regulators, and the effects on gene expression is therefore vital in elucidating the transcriptional control governing the regenerative process. To date, most studies have relied on transcriptomic studies to document gene expression changes in the regenerating liver^11–15^, while two others that focused on histone modifications^16,17^. However, these processes are downstream of chromatin reorganization and therefore do not capture the dynamic crosstalk of chromatin accessibility and transcriptional regulation. To identify transcriptomic changes specific to repopulating hepatocytes, we previously employed the translating ribosome affinity purification (TRAP)^18^ method followed by high-throughput RNA-sequencing (TRAP-seq) that coexpresses the GFP-tagged ribosomal protein, GFP-L10A, with FAH followed by affinity purification with anti-GFP antibodies to isolate translating mRNAs only from repopulating hepatocytes^15^. To discern the dynamic chromatin patterns that underlie liver repopulation, we now implement the ‘isolation of nuclei tagged in specific cell types’ (INTACT)^19^ method to isolate nuclei only from repopulating hepatocytes. This is achieved by expressing the GFP-tagged nuclear envelope protein SUN1-GFP with FAH in *Fah*^−/−^ mice, followed by sorting of GFP-positive nuclei from repopulating hepatocytes and ATAC-seq^20^. We identify promoter accessibility changes corresponding to upregulation of cell cycle pathways and downregulation of metabolic pathways, corroborating previous gene expression studies^12,15^. Integrative expression level and chromatin accessibility analysis suggests that gene activation is primarily associated with increased promoter accessibility, while inactivation is correlated with closure of selected promoters and enhancers. We propose a model in which a more accessible promoter allows increased transcription factor binding and gene activation, whereas decreased enhancer accessibility prevents binding of hepatocyte-enriched DNA binding proteins followed by inhibition of liver function genes so that the repopulating liver assumes a less differentiated state to promote cell growth and proliferation.

## RESULTS

### Adaptation of INTACT in the *Fah*^−/−^ model allows for isolation of repopulating hepatocyte nuclei

Liver cells in humans and mice rarely undergo division in homeostatic conditions^2^. However, with injury and repopulation, hepatocytes become facultative stem cells and divide to replenish liver mass and restore liver function^2^. We hypothesized that this change from quiescence to replication is accompanied by substantial and specific changes to chromatin accessibility. To analyze the chromatin specific to repopulating hepatocytes, we adapted the INTACT^19^ method in the *Fah*^−/−^ model to label hepatocytes with the GFP-tagged nuclear envelope, SUN1-GFP, and performed fluorescence-activated cell sorting (FACS) to isolate nuclei from whole liver at selected time points (Figure 1). The SUN1-GFP fragment was subcloned into a FAH expression plasmid^7^ so that all repopulating hepatocytes express GFP on the nuclear envelope. Following hydrodynamic injection of the FAH-SUN1-GFP plasmid into *Fah*^−/−^ mice, NTBC was removed and liver repopulation was allowed to proceed for one or four weeks (Figure 1A). As a control for healthy, quiescent hepatocytes, *Rosa*^LSL-SUN1-GFP^ transgenic mice^19^ were injected with AAV8-TBG-Cre^21^ to label all hepatocytes. Nuclei were isolated at the selected time points and FACS-sorted with an anti-GFP antibody to purify nuclei from repopulating hepatocytes exclusively (Figure 1B). ATAC-seq^20^ was then performed on the sorted nuclei to profile the changes in the chromatin regulatory landscape that occur during liver repopulation.

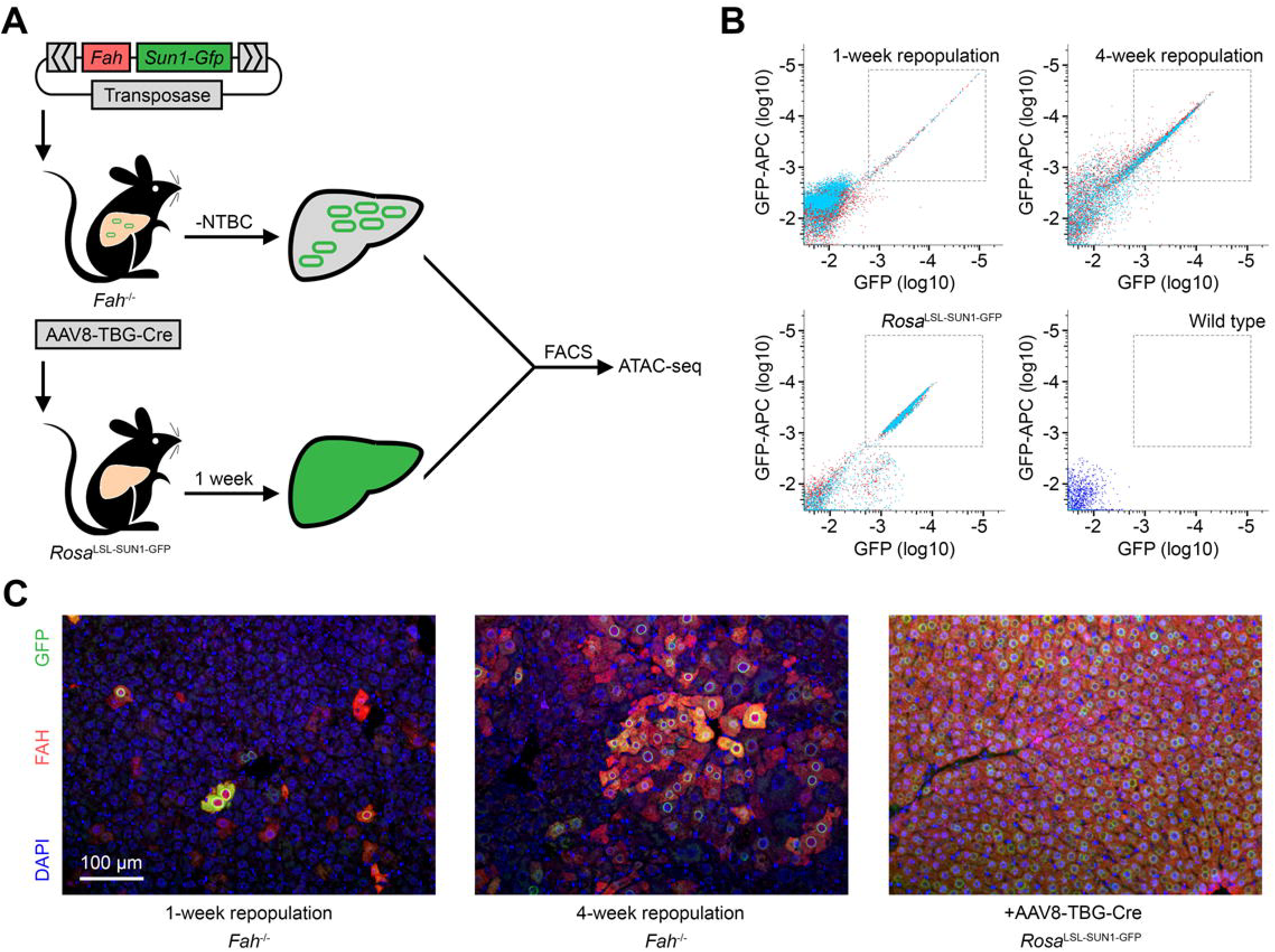

Immunofluorescence labeling demonstrated expression of GFP-tagged nuclear envelopes in FAH-positive cells (Figure 1C), illustrating the specificity of using SUN1-GFP^+^ nuclei as a marker to identify repopulating hepatocytes. Interestingly, FAH and GFP signal intensity were not homogeneous across all replicating cells, possibly due to the different copy numbers of plasmids taken in after hydrodynamic tail-vein injection of the SUN1-GFP construct^22^. In addition, since the *Sleeping Beauty* transposon system displays little insertion site preference^23^, the loci in which the DNA fragments are integrated can affect expression levels of FAH and SUN1-GFP^24^.

### ATAC-seq detects differentially accessible chromatin regions

All ATAC-seq libraries were sequenced to ~100 million reads to ensure ample coverage across the genome followed by quality assessment to verify the robustness of the data (Table 1). We observed consistent ATAC-seq signals across various loci such as the *Alb* gene, which showed a progressive decrease in accessibility at the enhancer region during repopulation (Figure 2A). To identify differentially accessible chromatin regions, fragments below 150 bp, termed ‘nucleosome-free reads’, were used for peak calling. We identified 16,043 differentially accessible regions between quiescent and repopulating hepatocytes (Figure 2B, Table 2), of which 1,244 displayed increased accessibility in 1-week and 1,266 increased accessibility in 4-week repopulating hepatocytes, while 2,058 regions showed decreased accessibility in week 1 and 2,036 decreased in week 4. Hierarchical clustering of the differentially accessible sites showed a clear separation of repopulating and quiescent hepatocytes (Figure 2C), corroborating with previous transcriptome studies that 1-week and 4-week repopulating hepatocytes have a similar expression profile distinct from quiescent hepatocytes^15^. Replicates also clustered within the same condition, illustrating the reproducibility between biological replicates. Comparing accessibility regulated in the same direction in both time points (‘congruent’), 1,241 peaks were congruently increased and 2,033 congruently decreased (Figure 2B). Of note, only 28 regions exhibit accessibility changes in opposite directions in week 1 and week 4 (‘incongruent’), reflecting the similarity in the chromatin profile between the two repopulation time points.

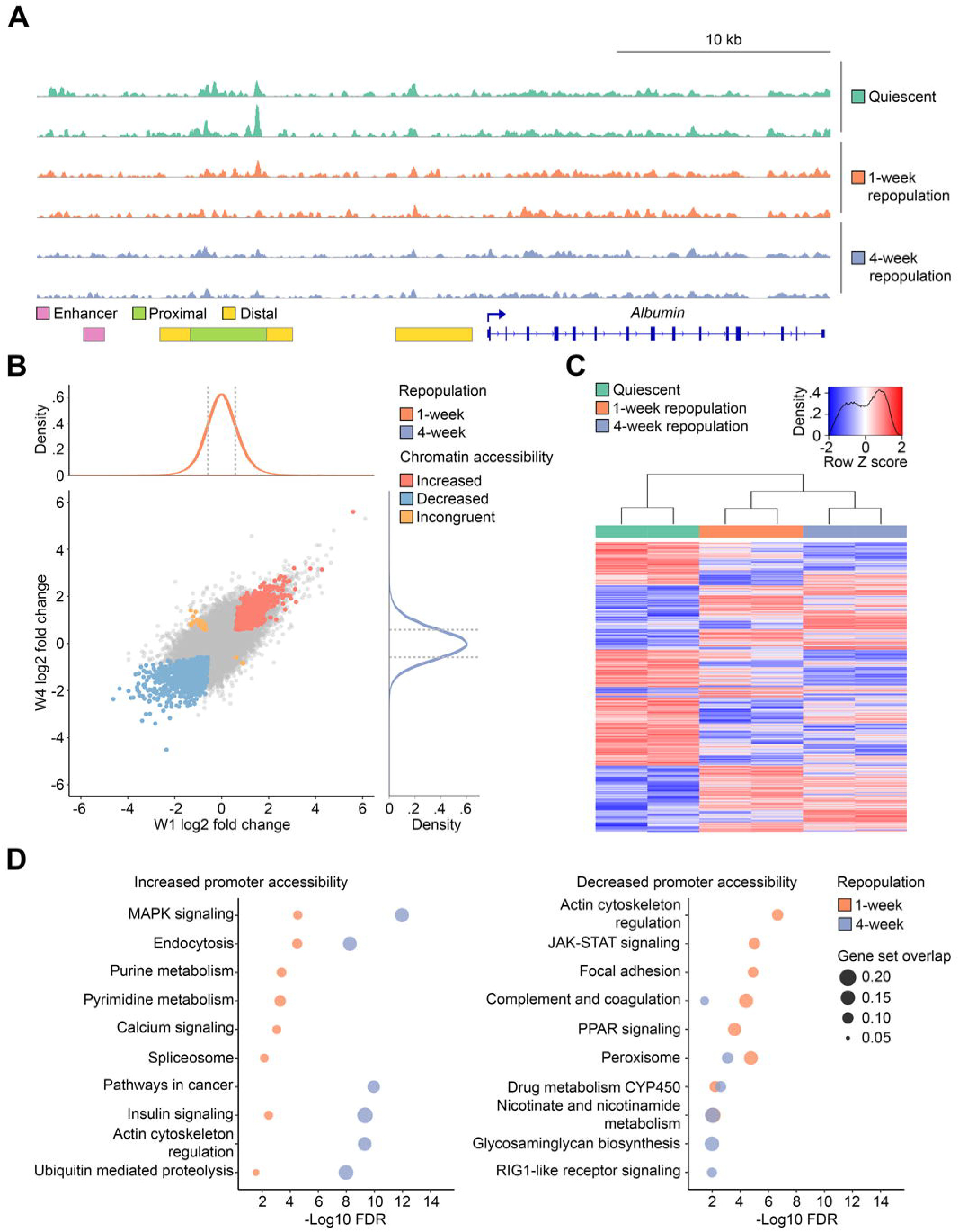

Next, we focused on differentially accessible promoter elements. Differential ATAC-seq regions within 1 kb up- and downstream of the transcription start sites (TSS) were determined and KEGG pathway^25^ analysis was performed (Figure 2D). As expected, pathways involved in cell growth and proliferation were enriched among the genes with increased accessibility in the promoter regions during repopulation, including MAPK signaling^26^ and cancer pathways. Interestingly, purine and pyrimidine metabolism are only enriched in genes with increased promoter accessibility at week 1 but not at week 4, suggesting early activation of DNA synthesis immediately after liver injury in early stages of repopulation. This observation is consistent with previous comparison of the *Fah*^−/−^ and partial hepatectomy (PHx) models showing that the transcriptome of 1-week repopulating hepatocytes in the *Fah*^−/−^ mouse is closest to that of 36 and 48 h post PHx^15^, at which the highest rate of DNA synthesis occurs in this model^27^. On the other hand, genes involved in mature hepatocyte functions such as complement and coagulation and metabolic pathways had significantly decreased promoter accessibility at both regeneration time points. Our pathway enrichment analysis substantiates prior studies of gene expression profiles and extends the findings to chromatin accessibility in that proliferation pathways are activated while liver functions are inhibited during repopulation^12,15^.

### Integration of chromatin accessibility and gene activity infers regulatory mechanisms

To evaluate the association of chromatin landscape and gene expression, we utilized our prior TRAP-seq study^15^ as a dataset of transcriptomic changes in repopulating hepatocytes. Genes with ATAC-seq signals and TRAP-seq reads that changed in the same direction at the same time point were identified as ‘concordant genes’ (Figure 3A, Table 3). We observed significant overlap of the concordant genes with ATAC-seq and TRAP-seq (p<1E-16 for all 1-week concordant genes and 4-week concordantly activated genes. p=0.03 for 4-week concordantly inhibited genes), while there was no significant overlap of genes with increased expression in 1 week and decreased chromatin accessibility at 4 week (p=0.39). The concordant target analysis indicates that changes to chromatin accessibility correspond to gene expression levels at specific time points. KEGG pathway^25^ analysis suggested enrichment of cell growth and replication in the week 1 concordantly activated genes, and overrepresentation of biosynthesis and metabolism in both week 1 and week 4 concordantly inhibited genes (Figures 3B, C). In addition, pathway enrichment supported previous observations that activation of the glutathione metabolic network is essential for reactive oxygen species removal after partial hepatectomy or recovery following toxic liver injury^15,28,29^. We conclude that changes to the chromatin structure underlie the upregulation of genes involved in proliferation and downregulation of genes associated with metabolic processes.

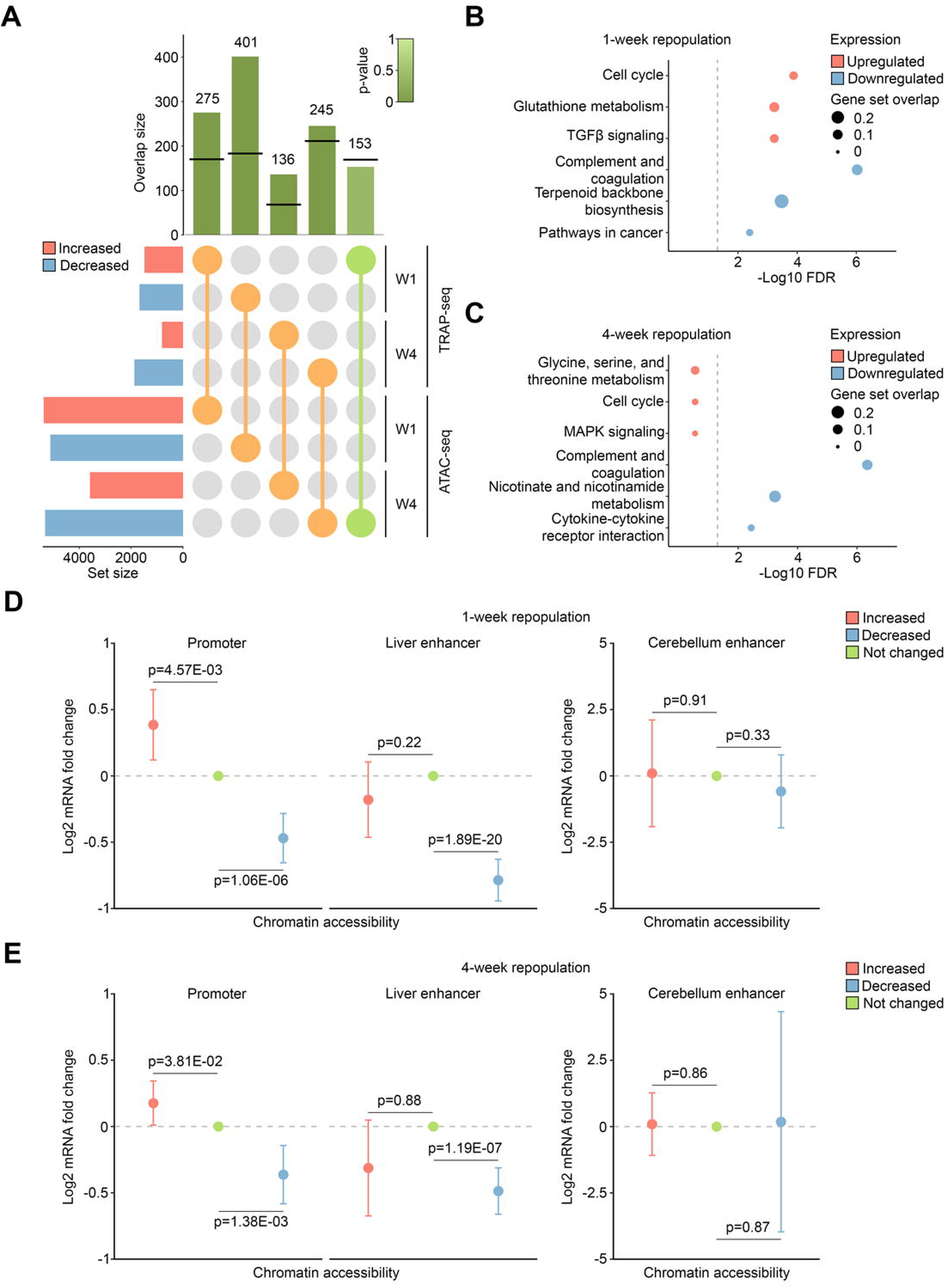

Next, we sought to investigate co-regulatory networks of functional regions and gene activity in repopulating hepatocytes. All ATAC-seq peaks identified were first separated into increased, decreased, or unchanged accessibility with a cutoff of absolute fold change ≥1.5 and false discovery rate (FDR) ≤0.05, followed by subdivision into regulatory regions of promoters, liver-specific enhancers, or cerebellum-specific enhancers as a negative control^30^. Promoter peaks were annotated to the nearest genes and the corresponding transcript levels at the same time point were extracted from TRAP-seq data^15^. We then compared the differential gene expression in the differentially accessible promoters to that in the unchanged promoters (Figures 3D, E). The normalized log_2_ fold change was positive (p=7.47E-03 in week 1 and 3.81E-02 in week 4) with increased and negative (p=1.06E-06 in week 1 and 1.38E-03 in week 4) with decreased promoter accessibility at both time points, demonstrating a significant association of promoter openness and transcriptional activity. Differentially accessible liver enhancer peaks were similarly categorized, putative enhancer-regulated genes extrapolated^30^, corresponding target gene expression extracted^15^, and the transcript level changes compared to those of genes with unchanged enhancer accessibility. Interestingly, decreased liver enhancer accessibility was highly correlated with decreased gene activity (p=1.89E-20 in week 1 and 1.19E-07 in week 4), while no significant expression changes (p=0.22 in week 1 and 0.88 in week 4) were associated with increased enhancer openness. The cerebellum enhancers exhibited no significant correlation with the changes in transcript levels and chromatin accessibility in the repopulating liver, as expected (Figures 3D, E, right). Our integrated ATAC-seq and TRAP-seq analysis reveals that gene activation is regulated by increased promoter accessibility, presumably allowing recruitment of transcriptional activators and RNA polymerase II to the TSS, whereas gene inhibition may be governed by both decreased promoter and enhancer openness, preventing long-range enhancer-promoter interactions^31^.

### Differential chromatin accessibility predicts transcription factor involved in liver repopulation

Dynamic coordination of chromatin structure and transcription factors is required to fine-tune gene expression. Chromatin organization influences access of the transcriptional apparatus by regulating binding sequence accessibility^32^ and transcription factor binding stability^33^; conversely, transcription factors affect access of remodelers to the chromatin^32^ and histones^34^. To identify DNA binding transcription factors that connect differential chromatin accessibility and gene expression, we carried out *de novo* motif profiling at differentially accessible promoters and liver enhancers^30^.

We found enrichment of the ETS transcription factor ELK1 motif in promoters with increased accessibility in both 1-week (FDR=1E-76) and 4-week (FDR=1E-41) repopulating hepatocytes (Figure 4A, B). ELK1 binds to the serum response element upon MAPK phosphorylation^35^ to activate immediate early genes such as *Fos* and components of the basal transcriptional machinery^36^. Furthermore, ELK1 supports cell cycle entry during liver regeneration as *Elk1*^−/−^ mice show reduced hepatocyte proliferation after PHx^37^. We postulate that promoters became more accessible after acute liver injury to permit increased ELK1 occupancy, enabling hepatocyte repopulation.

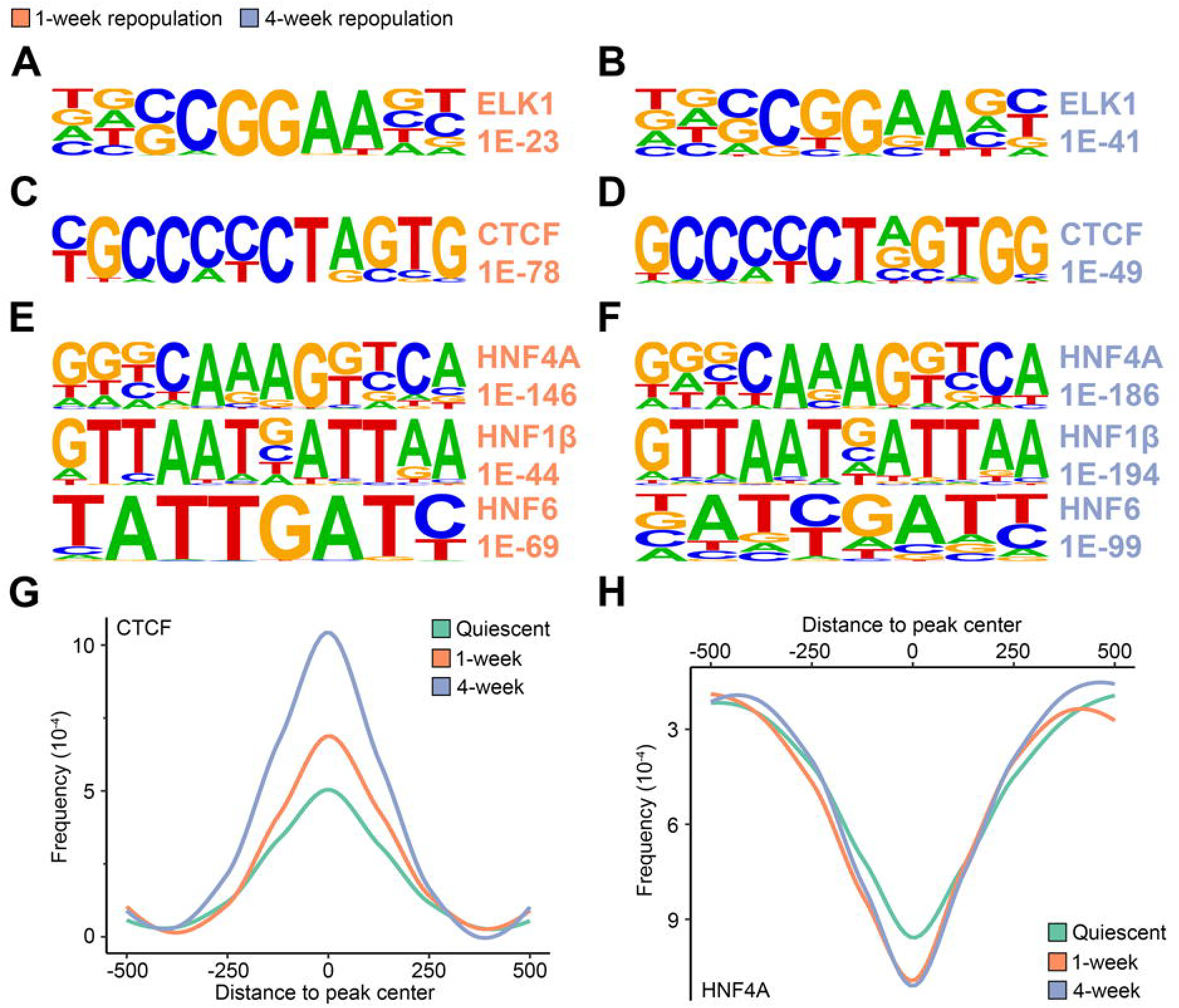

Among the regions with increased accessibility during liver repopulation, surprisingly, the CTCF motif was highly enriched (FDR=1E-78 in week 1 and 1E-49 in week 4) (Figures 4C, D). CTCF plays numerous roles in transcriptional regulation to function as a transcriptional activator^38^, repressor^39^, insulator to block enhancer-promoter interactions^40^, chromatin structure organizer to form topologically-associated domains^41^ modulator of long-range chromatin looping^42^, and even mediator of local RNA polymerase II pausing to regulate alternative exon usage^43^. However, the function of CTCF in liver regeneration has not been studied to date.

In addition, we found the HNF4α binding motif to be significantly associated with liver enhancers with decreased accessibility during liver regeneration (FDR=1E-146 in week 1 and 1E-186 in week 4) (Figures 4E, F). HNF4α is a master regulator atop the transcriptional cascade of hepatocyte differentiation^44,45^ and a crucial factor that maintains hepatocytes in the differentiated state^46^. Importantly, HNF4α suppresses liver proliferation, as mice with conditional deletion of *Hnf4a* demonstrate increased hepatocyte BrdU incorporation and Ki67 expression^47^. HNF4α also directly inhibits cell growth and replication pathways, as illustrated by upregulation of cell cycle and proliferation genes upon acute HNF4α loss^47,48^. Moreover, motifs of other liver-enriched transcription factors were also overrepresented at enhancers that became less accessible in repopulating hepatocytes, including hepatocyte nuclear factor 1β (HNF1β) and hepatocyte nuclear factor 6 (HNF6)^49^ (Figure s4E, F)^49^. We examined the locations for CTCF and HNF4α motifs within regions of dynamic chromatin accessibility and found that they are present in the center of these regions with CTCF at those with increased, and HNF4α at those with decreased accessibility (Figure 4G, H).

In summary, *de novo* motif analysis of the ATAC-seq footprints suggests increased occupancy of ELK1 and CTCF at chromatin regions that become more accessible, and decreased binding of liver-enriched transcription factors at liver enhancers that become less accessible during repopulation.

## HNF4α occupancy is decreased in liver-specific enhancers during repopulation

We postulated that decreased HNF4α binding allows repopulating hepatocytes to assume a less differentiated and pro-proliferative state and carried out ChIP-seq on quiescent and 4-week repopulating livers to examine genome-wide HNF4α occupancy during the repopulation process. We observed 508 peaks with decreased and only 14 peaks with increased occupancy in repopulating livers (Figure 5A, Table 4). Remarkably, 42% (214) of lost HNF4α occupancy occurred within previously-defined liver enhancers^30^, while 23% (119) fell into distal intergenic regions, and 10% (52) were within 1 kb up- and downstream of TSS (‘promoter’) (Figure 6B). These data corroborate the differentially accessible chromatin footprint analysis that had identified the HNF4α motif at enhancers with decreased accessibility in repopulating hepatocytes (Figure 4H).

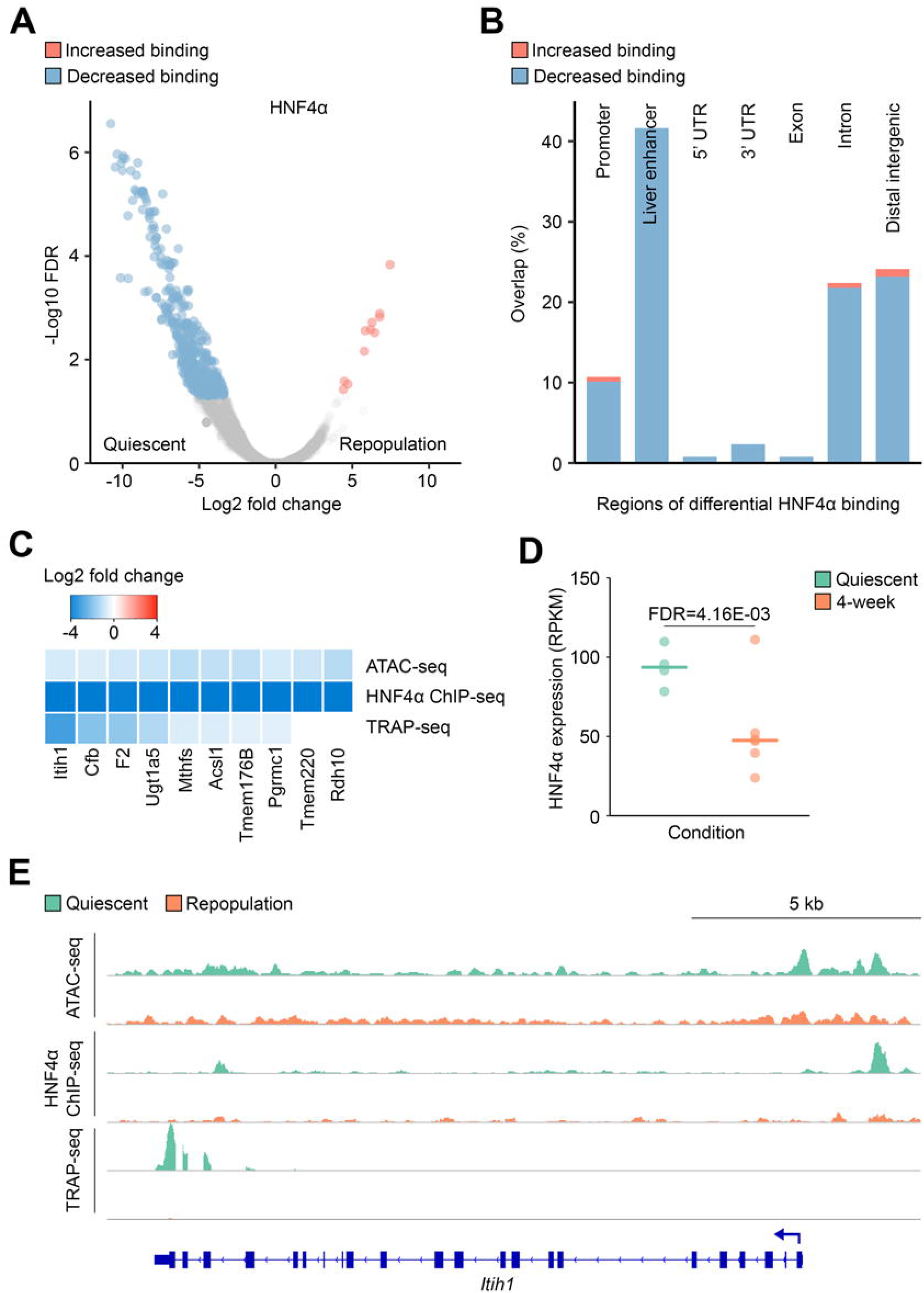

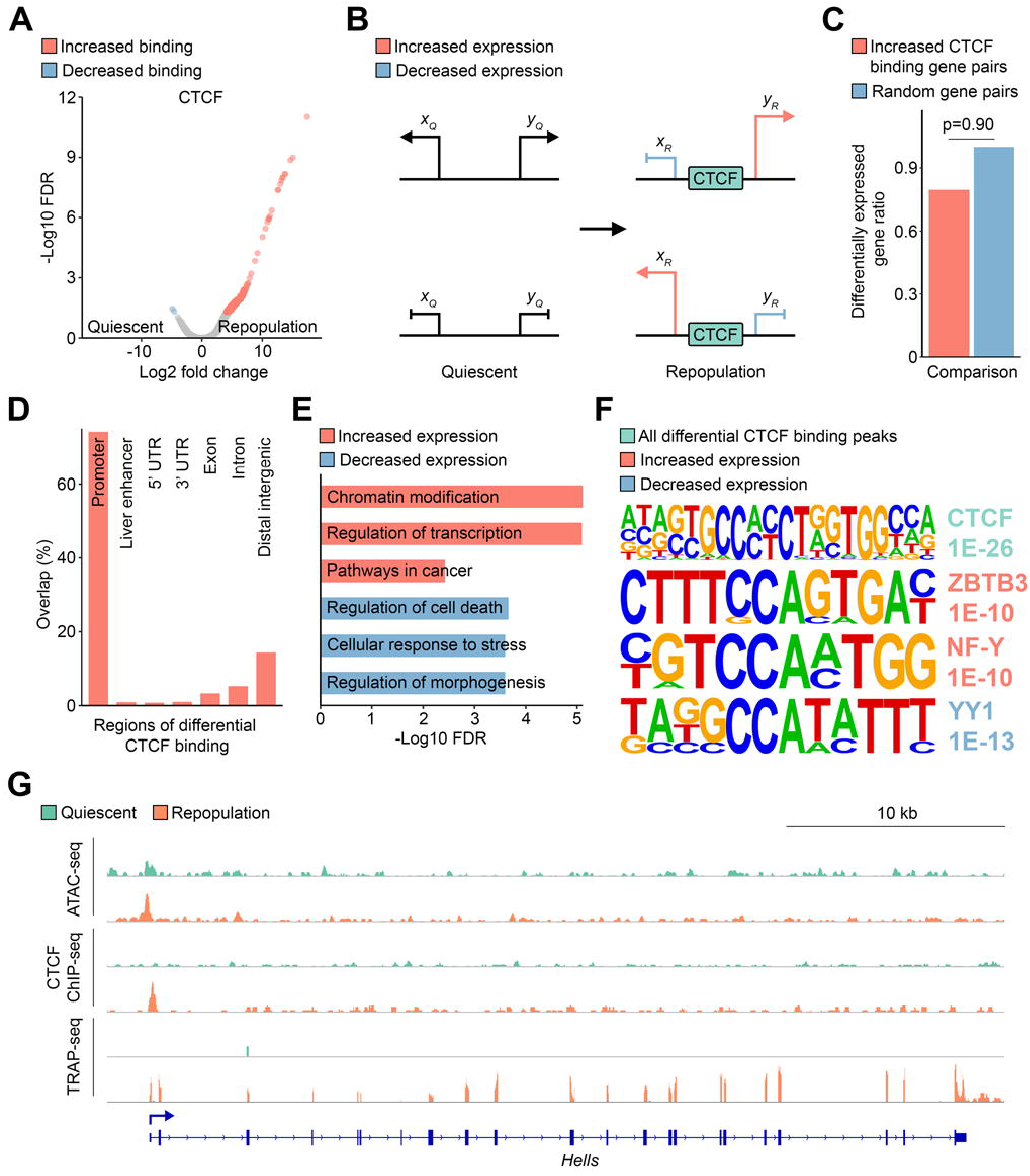

Next, we integrated ATAC-seq, ChIP-seq, and TRAP-seq datasets^15^, and identified hepatocyte-enriched genes crucial for establishing liver functions including complement and coagulation (*Cfb*, *F2*), biosynthesis (*Itih1*, *Acsl1*, *Pgrmc1*), and metabolism (*Ugt1a5*, *Mthfs*, *Rdh10*)^50^ as correlated with decreased HNF4α enhancer occupancy during regeneration (Figure 5C, E). To explore the mechanism responsible for decreased HNF4α occupancy during liver repopulation, we next turned to the TRAP-seq dataset^15^ to inspect *Hnf4a* expression levels in quiescent and replicating hepatocytes. Remarkably, we found a 50% reduction of *Hnf4a* transcripts in 4-week repopulating hepatocytes (FDR=4.16E-3) compared to the quiescent liver (Figure 5D). Taken together, these results implicate decreased chromatin accessibility and reduced *Hnf4a* expression as contributors to suppression of hepatocyte-specific genes and downregulation of liver biosynthetic functions during repopulation.

### CTCF promoter occupancy is increased in the repopulating liver

In order to extend the computational finding of enriched CTCF motif at promoters with increased accessibility, we performed ChIP-seq in quiescent and 4-week repopulating livers. CTCF occupancy was increased at 1,382 sites in the repopulating liver, while only 2 peaks showed decreased binding (Figure 6A, Table 5). To characterize the role of increased CTCF occupancy during liver repopulation, we first evaluated its potential insulator function by calculating an ‘insulator strength score’^51^ at all gained binding sites. Genomic regions with increased CTCF occupancy with divergent flanking promoters within 50 kb were identified and the normalized expression levels corresponding to the genes extracted from our TRAP-seq data^15^. Surprisingly, gene pairs with increased CTCF binding were not significantly more enriched for differential gene expression than random gene pairs (p=0.9), suggesting that CTCF is unlikely to act as a differential expression insulator during liver repopulation.

Remarkably, the vast majority (1,026, 74%) of the gained CTCF peaks fell within 1 kb up- and downstream of the TSS (‘promoter’) (Figure 6D). To examine the targets of increased CTCF occupancy, all differentially bound peaks were annotated to the nearest genes and their corresponding expression changes were obtained from our TRAP-seq dataset^15,25,52^. We found 545 (39%) peaks associated with activation of genes in cell growth and proliferation pathways such as chromatin modification, transcription regulation, and cancer (Figure 6E), while 656 (47%) sites with increased CTCF binding were associated with inhibition of genes in cell death regulation, stress response, and morphogenesis. Together, our network analysis suggests a diverse role for CTCF in transcriptional regulation in which increased CTCF occupancy could support hepatocyte replication and prevent cell death during liver repopulation, possibly by enabling binding of both activating and repressing cofactors.

Indeed, CTCF is known to exhibit divergent roles in activating and repressing transcription by recruiting various protein partners in a context-dependent manner^53^. To identify these cofactors, we performed motif analysis for the regions differentially bound by CTCF (Figure 6F). As expected, the CTCF motif was highly enriched (FDR=1E-26) at all differential binding sites, confirming the specificity of the anti-CTCF antibody for immunoprecipitation. At sites where CTCF binding corresponded to gene activation, we observed significant enrichment for ZBTB3 (FDR=1E-10) and NF-Y (FDR=1E-10) binding motifs (Figure 6F). ZBTB3 is considered a likely factor binding 5’ of CTCF due to its frequent enrichment ~10 bp upstream of CTCF motifs in the human genome^54^. Furthermore, expression of ZBTB3 is induced by accumulation of reactive oxygen species to promote cancer cell growth and prevent apoptosis via activation of antioxidant gene expression in cell lines^55^. Whether CTCF directly interacts with or indirectly recruits ZBTB3 is as yet unclear, but the proteins are likely to interact based on their close proximity at promoters to enhance transcription in repopulating hepatocytes. The transcription factor NF-Y binds to the CCAAT box present at ~30% of the promoters^56^ and is required for cell cycle progression, DNA synthesis, and proliferation in mouse embryonic fibroblasts^57^. Additionally, reconstituted *in vitro* transcription reactions demonstrated that binding of NF-Y disrupts nucleosome structure at promoters containing the NF-Y recognition sequence^58^. Recruitment of NF-Y could therefore induce local nucleosome repositioning to allow increased accessibility of the transcriptional apparatus to activate gene expression.

On the other hand, the Yin Yang 1 (YY1) binding motif was enriched (FDR=1E-13) at sites where increased CTCF occupancy corresponded with decreased gene expression (Figure 6F). YY1 regulates embryogenesis, cell differentiation, and tumorigenesis^59,60^, as well as enhancer-promoter interactions analogous to long-range chromatin looping mediated by CTCF^61^. YY1 functions as a transcriptional repressor via recruitment of the polycomb repressor complex, resulting in trimethylation of histone H3 lysine 27^62,63^. It is also a cofactor of CTCF in regulating X chromosome inactivation, although the exact mechanism remains unclear^64^. Given these observations, it is likely that direct or indirect co-binding of CTCF and YY1 at promoters induces transcriptional repression or disrupts enhancer to promoter interactions to downregulate target genes. These results suggest that increased chromatin accessibility correlates with increased CTCF occupancy to recruit coactivators or corepressors to fine-tune target gene expression to induce cell replication, and prevent cell apoptosis during liver repopulation (Figure 6G).

### Liver regeneration is accompanied by nucleosome remodeling

Most eukaryotic DNA is packaged around histone protein octamers into nucleosomes to regulate chromatin organization and transcriptional control. Nucleosome properties such as positioning and turn-over rates can affect binding of transcription factors and access of the transcriptional machinery^65^. The nucleosome landscape adjacent to the TSS is of particular interest, as nucleosomes adopt a specific phasing pattern immediately up- and downstream^66^. Hence, nucleosome organization could act as an additional layer of transcriptional regulation in the repopulating hepatocytes.

We inferred nucleosome positioning from nucleosome-containing sequences by extracting ATAC-seq reads longer than 150 bp (Figure 7A). Nucleosomes surrounding the TSS were defined as ‘-1 nucleosomes’ within 350 bp upstream and ‘+1 nucleosomes’ within 250 bp downstream, and the distance between the +1 to −1 nucleosomes was defined as the ‘nucleosome-free region’. When compared to quiescent hepatocytes, there was a median downstream shift of 9 bp in 1-week (p=2.60E-13) and an upstream shift of 19 bp in 4-week (p<1E-15) repopulating hepatocytes for the −1 nucleosomes, while there was no significant shift in +1 nucleosome positioning (Figure 7B, Table 6). As a result, there was a global increase of promoter openness in 4-week repopulating hepatocytes as the distance between +1 to −1 nucleosomes increased, while the nucleosome-free region was more closed in 1-week regenerating liver compared to the quiescent state.

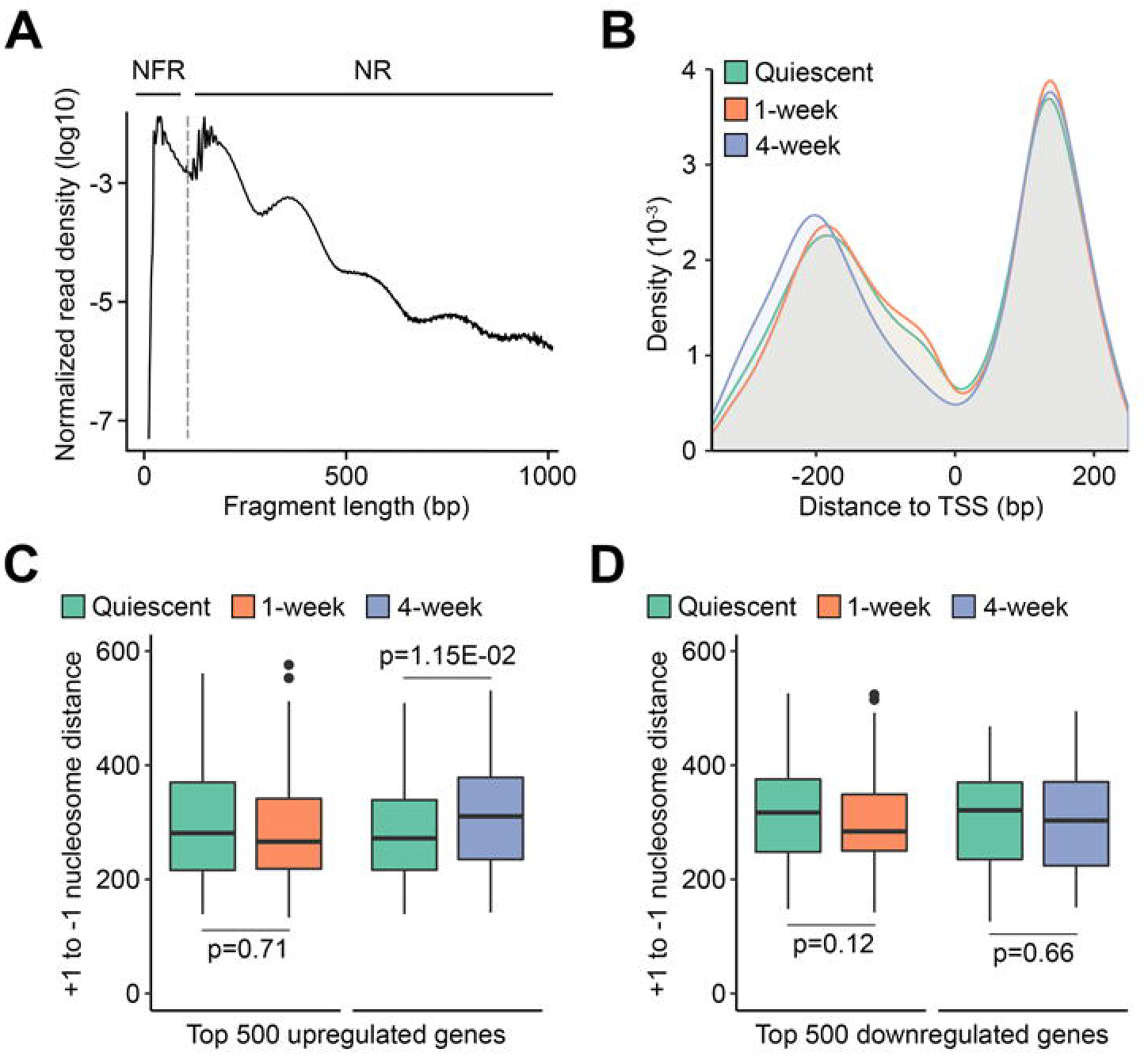

To evaluate the association of TSS accessibility and gene expression, we extracted the top 500 up- and downregulated genes in repopulation^15^ and calculated the change in the length of the nucleosome-free region between quiescent and regenerating hepatocytes as a surrogate for differential TSS accessibility. We only observed a significant increase (p=1.15E-2) of +1 to −1 nucleosome distance in genes activated in week 4 when compared to quiescent hepatocytes, while no significant change in nucleosome-free region was present in genes upregulated in week 1 or genes downregulated in week 1 and week 4 (FIgure 7C, D). It is likely that eviction or repositioning of the −1 nucleosomes could expose transcription factor binding sequences and allow access of the transcriptional machinery to the TATA box for gene activation in regenerating hepatocytes^67^. Altogether, analysis of the nucleosome structure implies nucleosome reorganization could affect gene activation but not inhibition during liver repopulation.

## DISCUSSION

Gene regulation is tightly controlled by a complex network integrating transcription factor binding and transcriptional apparatus assembly, chromatin structure, epigenetic modifications, and even intra- and interchromosomal interactions^9,10^. In this study, we investigated the association of chromatin accessibility, nucleosome properties, transcription factor occupancy, and gene expression^15^ to delineate the multidimensional framework of transcriptional regulation in the repopulating liver. By implementing the INTACT method^19^ to express SUN1-GFP in the *Fah*^−/−^ model, we successfully performed cell type-specific isolation of only repopulating hepatocyte nuclei followed by ATAC-seq to identify changes of the chromatin landscape (Figures 1, 2). Integration of TRAP-seq^15^ with ATAC-seq determined that gene activation corresponds with increased promoter openness, while gene inhibition is linked to decreased promoter and enhancer accessibility (Figure 3C). We also corroborated previous findings that cell cycle, DNA synthesis, proliferation, and glutathione metabolism are activated whereas complement and coagulation, biosynthesis, and metabolic pathways are inhibited during liver repopulation (Figures 2D and 3B, C)^12,15^. In addition, *de novo* footprint analysis identified enrichment of CTCF and HNF4α motifs in regions with increased and decreased accessibility in repopulating hepatocytes, respectively (Figure 4). We further validated differential occupancy of both factors in the repopulating liver with ChIP-seq and observed decreased HNF4α binding at liver enhancers^30^ (Figure 5) and increased CTCF binding at promoters (Figure 6). Integrated ATAC-seq, ChIP-seq, and TRAP-seq analysis suggests that CTCF recruits cofactors to activate genes involved in chromatin organization and replication and inhibit genes in the regulation of cell death (Figure 6E-G). On the other hand, loss of HNF4α occupancy at liver enhancers decreases expression of hepatocyte-enriched genes crucial in establishing liver homeostasis and function (Figure 5C-E).

In general, 40% of CTCF binding sites occur in intergenic regions distant to TSS, while 35% of CTCF sites are found in promoters^30,41^. Interestingly, the vast majority (75%) of sites with increased CTCF occupancy are located within promoters in the repopulating liver (Figure 6D). In fact, CTCF can function as a direct transcriptional repressor at the *c-myc* promoter^68^ and as an activator of the amyloid precursor protein promoter^69^, strengthening the notion that CTCF plays a more localized role as a transcriptional regulator in the repopulating liver via recruitment of cofactors. Nonetheless, the multitude of CTCF functions warrants further investigation to understand its contribution to mediating chromatin structure and organization in the context of liver repopulation. Specifically, CTCF also acts as an insulator to block enhancer-promoter interactions^40^, a factor that promotes long-range chromatin looping^42^, and a TAD boundary protein that defines expression domains for tight transcriptional control^41^. Future experiments to detect changes in chromatin interactions via chromosome conformation capture^70^ would be valuable in directly determining whether differential CTCF occupancy affects three-dimensional chromatin organization during liver repopulation.

The mechanisms of increased CTCF and decreased HNF4α binding in the repopulating liver are also not fully understood. In the current study, we infer that a more open chromatin state at specific promoters correlates with accessibility of CTCF to its binding sites; however, we have no way of knowing causality. Previous work found that enrichment of thymidine (T) at the 18^th^ position in the CTCF motif reduces its affinity, where low-affinity sites are more sensitive to CTCF binding gain and loss during mouse embryonic stem cell differentiation^51^. Additionally, it is likely that changes in DNA methylation influence differential CTCF occupancy, as methylated CpGs in the CTCF recognition site can prevent its binding^71,72^. Demethylation at specific promoter regions could therefore increase CTCF occupancy during liver repopulation. In the case of reduced HNF4α occupancy at liver-specific enhancers in the regenerating liver, part of this effect can be explained by reduced expression of HNF4α itself. Furthermore, HNF4α could be regulated post-transcriptionally via phosphorylation by kinases such as protein kinase A and C, as well as AMP-activated protein kinase to decrease its DNA binding activity or nuclear localization^73^. Activation of the MAPK signaling pathway is also shown to inhibit *Hnf4a* expression via activation of the transcription factor c-Jun^73,74^. The fact that enrichment of DNA synthesis pathways are only observed in 1-week repopulating livers and that *Hnf4a* transcript level is unchanged in week 1 but reduced in week 4 hepatocytes strengthens the notion that activation of cell growth and proliferation occur early after the initiation of liver repopulation, followed by a later inhibition of *Hnf4a* transcription. Future studies using, for instance, targeted degradation of CTCF^75^ or HNF4α could be implemented to identify potential promoters and inhibitors of liver repopulation. Technologies such as cDNA^8^ or clustered regularly interspaced short palindromic repeats (CRISPR)^76,77^ screens could also be utilized to evaluate the effectors downstream of CTCF activation and HNF4α inhibition.

In summary, we propose the following model to explain the transcriptional adaptations that accompany liver repopulation (Figure 8): during hepatocyte replication, the promoters of selected genes become more open due to an increased distance between histones at +1 to −1, allowing increased sequence accessibility for CTCF, transcription factor recruitment, and transcriptional machinery assembly to activate genes that regulate cell cycle, DNA synthesis, and proliferation pathways. On the other hand, decreased enhancer accessibility in conjunction with suppression of *Hnf4a* expression evicts or prevents HNF4α binding, and possibly that of other hepatocyte nuclear factors, to liver enhancers, resulting in repression of hepatocyte metabolic and biosynthetic function genes.

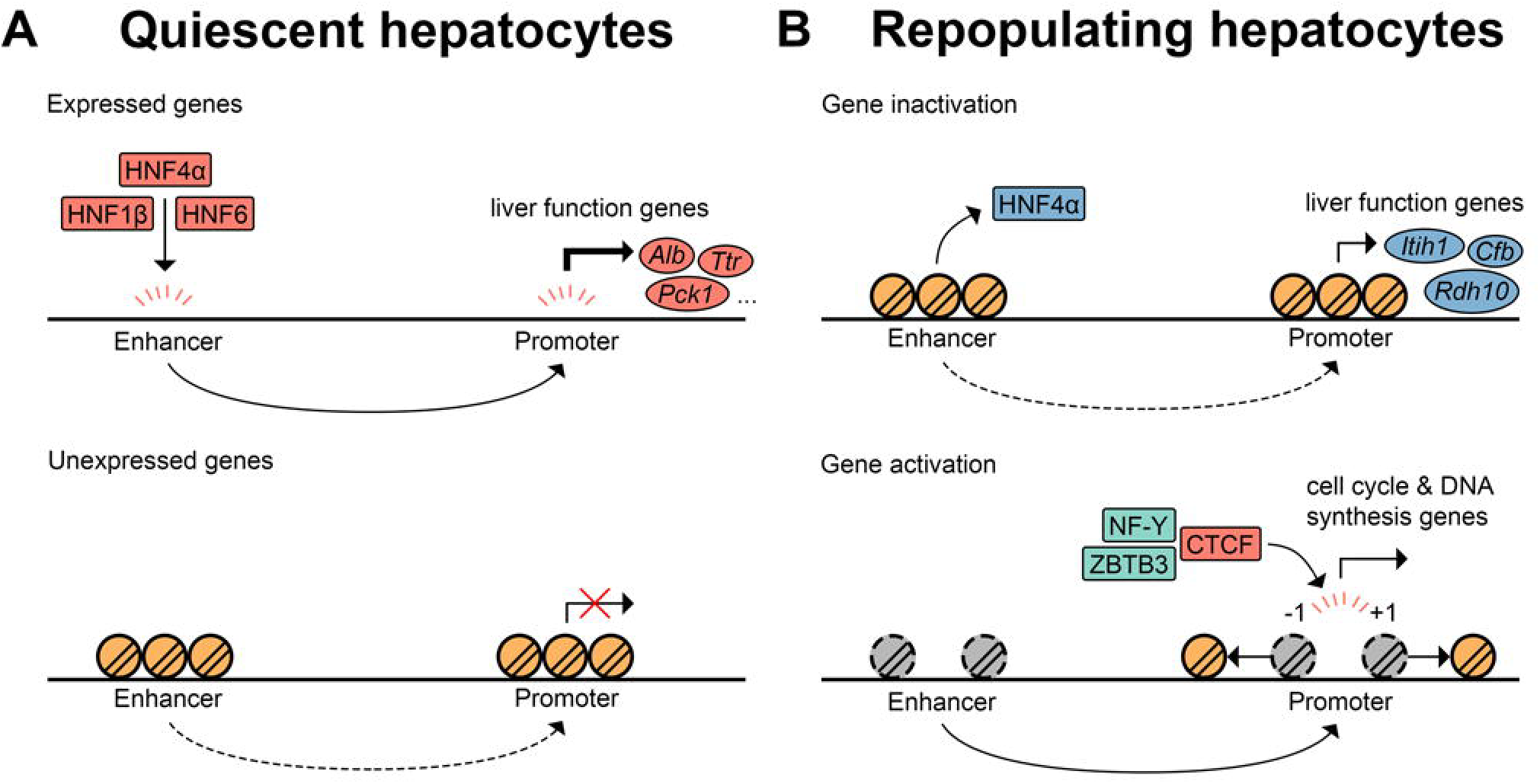

## METHODS

### Plasmid construction

The generation of the pKT2/Fah-Sun1-Gfp//SB plasmid was described previously^15^. The nuclear envelope SUN1-tagged GFP (SUN1-GFP) plasmid was a generous gift from Dr. Jeremy Nathans (Johns Hopkins University, Baltimore, MD, USA). We amplified the SUN1-GFP insert by PCR amplification with the primers MfeI-Sun1-F (GACTCAATTGGCGGCCGCACTACTGGCC) and BsiW1-Sun1-R (GCTACGTACGTTAACCGCTACTATTAAGATCCTCCTCGGATATTAACTTCTGC) and subcloned it into the vector pKT2/Fah-mCa//SB^7^ to construct pKT2/Fah-Sun1-Gfp//SB. This construct utilizes the *Sleeping Beauty* (SB) transposase for stable transgene integration into the genome. The plasmid was prepared with the GenElute HP Plasmid Maxiprep Kit (NA0310-1KT, MilliporeSigma) for endotoxin-free maxi-scale DNA extraction and purification.

### Mouse studies

*Fah*^−/−^ mice were maintained on 7.5 mg/l 2-(2-nitro-4-trifluoromethylbenzoyl)-1,3-cyclohexanedione (NTBC) (Swedish Orphan Biovitrum) in the drinking water. Hydrodynamic tail-vein injection^76^ of 10 μg of pKT2/Fah-Sun1-Gfp//SB was performed followed by NTBC withdrawal for 1 week (n=2) or 4 weeks (n=2) to induce liver repopulation^15^. The *Rosa*^LSL-Sun1-GFP^ mice^19,78^ were kindly provided by Dr. Jeremy Nathans (Johns Hopkins University, Baltimore, MD, USA) and were tail-vein injected with AAV8.TBG.PI.Cre.rBG (Penn Vector Core) at 1 × 10^11^ virus particles per mouse to ablate the loxP-stop-loxP cassette only in hepatocytes. Livers from these mice were harvested 1 week after viral injection and served as quiescent controls. All studies were performed in 8 to 12-week-old mice.

### Immunofluorescence staining

Liver lobes were isolated, fixed in 4% paraformaldehyde overnight at 4°C, embedded in paraffin, and sectioned. Tissue sections were deparaffinized with xylene and rehydrated with serial incubation of 100%, 95%, 80%, and 75% ethanol followed by PBS. Antigen retrieval was carried out in Tris/EDTA buffer (10mM Trix, 1mM EDTA, pH 9.2) in a pressure cooker (2100 Antigen Retriever, Aptum Biologics Ltd.) and cooled to room temperature. Slides were then blocked with blocking buffer (PBS, 1% BSA) for an hour followed by overnight incubation of antibodies in the blocking buffer at 4°C in a humidified chamber. Three washes of PBS were carried out the next day followed by incubation with secondary antibodies at room temperature for 2 hours. Goat anti-GFP antibody (ab6673, 1:300, Abcam) and rabbit anti-FAH antibody (ab81087, 1:600, Abcam) were used to label repopulating hepatocytes from *Fah*^−/−^ mice after one and four weeks of repopulation and all hepatocytes from *Rosa*^LSL-GFP-L10a^ mice injected with AAV8-TBG-Cre. DAPI (B1098, 1:10,000, BioVision) was used to label nuclei.

### Hepatocyte nuclei isolation

Liver was homogenized in 10 ml hypotonic buffer (10 mM Tris-HCL, pH 7.5, 2 mM MgCl_2_, 3 mM CaCl_2_) on ice. The homogenate was filtered with a 100 μm filter and sedimented at 400 g for 10 minutes at 4°C. 10 ml of hypotonic buffer with 10% glycerol was used to resuspend the pellet followed by dropwise addition of 10 ml cell lysis buffer (hypotonic buffer, 10% glycerol, 1% IGEPAL CA-630). The homogenate was incubated for 5 minutes on ice and sedimented at 600 g for 5 minutes at 4°C. Nuclei were washed with lysis buffer again and quantified in a hemocytometer. Isolated nuclei were labeled with an Alexa Fluor 647 anti-GFP antibody (338006, clone FM264G, 1:25, BioLegend, San Diego, CA) for 30 minutes and 2 μg/ml DAPI immediately prior to sorting. After gating for the DAPI-positive signal, nuclei double positive for GFP and AF647 were sorted with a BD FACSAria II, and only tetraploid hepatocyte nuclei were collected for further experiments.

### ATAC-seq library generation

ATAC-seq libraries were generated as previously described^20^. Briefly, transposition was performed on 25,000 sorted tetraploid nuclei at 37°C for 30 minutes followed by DNA purification with the MinElute Reaction Cleanup Kit (28206, QIAGEN). DNA fragments were PCR preamplified for 5 cycles initially, and 1/10 of the volume (5 μl) was removed for qPCR amplification for 20 cycles. A ‘R vs Cycle Number’ plot was generated and the number of cycles required to reach ⅓ of the maximum R determined for each sample. The preamplified ATAC-seq libraries were then amplified for the calculated additional cycles. Agencourt AMPure XP beads (A63881, Beckman Coulter) were used for size selection to generate the final libraries^79^. Library quality was assessed with an Agilent High Sensitivity DNA Bioanalyzer (5067-4626, Agilent Technologies), and quantity measured with KAPA Library Quantification Kits (KK4835, KAPA Biosystems)

### ATAC-seq peak calling

ATAC-seq libraries were paired-end sequenced on an Illumina HiSeq 4000 (Illumina, San Diego, CA, USA) with 50, 75, or 100 reads. Reads were then trimmed to 50 bp with Cutadapt^80^ and peaks called with the ATAC-Seq/DNase-Seq pipeline (https://github.com/kundajelab/atac_dnase_pipelines). Briefly, the trimmed fastq files were aligned to the mouse genome (mm10) with Bowtie2^81^ followed by removal of PCR duplicates and mitochondrial reads. Bam files of the same biological sample from various technical replicates were then merged with Samtools^82^ and duplicated reads removed. The filtered reads were shifted 5 bp for + strands and 4 bp for -strands to adjust for the transposase binding sites^20^. Nucleosome-free reads were identified with the R package ATACseqQC using a random forest classifier^83^ followed by peak calling with MACS2^84^. Artifact signals were then removed according to the mm10 empirical blacklist regions^85^. The irreproducible discovery rate (IDR) framework was used to compare all pairs of biological replicates to identify reproducible peaks that passed a threshold of 10% for all pairwise analyses. The conservative peak set for each sample was identified by selecting the longest peak list from all pairs that passed the 10% IDR cutoff.

### ATAC-seq peak quality assessment

To ensure the ATAC-seq peaks generated from the sorted nuclei are of high quality, The R package ATACseqQC^83^ was employed for assessment. We first visualized the insert size distribution to confirm the presence of distinct periodicity of ~175 bp associated with nucleosome patterning in all samples, indicating the DNA fragments are protected by integer multiples of nucleosomes^20^. The signal intensity of nucleosome-free reads and nucleosomal reads was also averaged across all TSS to examine evidence that no over-fragmentation was introduced during hepatocyte nuclei isolation, sorting, or ATAC-seq library preparation.

### ATAC-seq differential peak analysis

The R package ATACseqQC^83^ was used to split the aligned bam files into nucleosome-free reads and nucleosomal reads. The R package DiffBind^86^ was used to identify differential accessible peaks from the nucleosome-free reads. The overlapping regions from the ATAC-seq peak sets for each sample were identified and merged into non-overlapping regions. Read counts for each region were quantified with dba.count (score=DBA_SCORE_TMM_READS_FULL, fragmentSize=0, bScaleControl=F, filter=0, bRemoveDuplicates=F, bUseSummarizeOverlaps=T). Peaks identified in both biological replicates in the same conditions were used for differential analysis with dba.analyze (method=DBA_EDGER, bSubControl=F, bTagwise=T) in conjunction with edgeR^87^. Peaks with an absolute fold change ≥1.5 and FDR ≤0.05 were identified as significant differentially accessible regions.

### Integrative analysis of TRAP-seq and ATAC-seq data

To identify chromatin accessibility and gene expression that changed in the same direction at the same time point (‘concordant genes’), the differentially accessible peaks were first annotated to the nearest TSS with the R package ChIPseeker^88^. Genes with differential expression during liver repopulation were obtained from a previous study that utilized translating-ribosome affinity purification followed by RNA-sequencing (TRAP-seq)^15^. The concordant ATAC-seq peaks and TRAP-seq genes were identified and the expected overlap and significance was calculated with a hypergeometric test. To evaluate the association of chromatin accessibility and gene expression changes, all chromatin regions were stratified into regions with increased, decreased, or unchanged accessibility, with the cutoff of an absolute fold change ≥1.5 and FDR ≤0.05. For promoter accessibility and gene activity association analysis, regions within 1 kb up- and downstream of the TSS were identified and annotated to the nearest genes with the R package ChIPseeker^88^. The corresponding expression change at the same time point was extracted from TRAP-seq^15^ and normalized by subtracting the mean log_2_ fold change of the unchanged from the increased and decreased chromatin accessibility groups. The normalized expression fold change of the nearest genes in the differentially accessible promoters was compared to that in the unchanged accessibility promoters with a one-sample t test. For enhancer accessibility and gene expression association studies, liver- and cerebellum-specific enhancers and their putative targets were obtained^30^. Gene expression fold changes were normalized as described above, and the normalized gene expression fold-change of the enhancer target genes in the differentially accessible enhancers was compared to that in the unchanged accessibility enhancers with a one-sample t test.

### ChIP-seq library generation

100 mg of quiescent (n=2) and repopulating (n=2) liver tissue was finely chopped with a razor blade and cross-linked in 1% formaldehyde for 10 minutes followed by addition of 2.5 M glycine and incubation for 5 minutes at room temperature. Tissues were sedimented, washed with cold PBS, and Dounce-homogenized in cold ChIP cell lysis buffer (10 mM Tris-HCl pH 8.0, 10 mM NaCl, 3 mM MgCl2, 0.5% IGEPAL CA-630, protease inhibitor) on ice. After incubation at 4°C for 5 minutes, nuclei were pelleted and resuspended in nuclear lysis buffer (50 mM Tris-HCl pH 8.1, 1% SDS, 5 mM EDTA, protease inhibitor). Nuclei were sonicated with a Bioruptor (Diagenode) for 2 rounds of 7.5 minutes each. 10 μg of sheared DNA was incubated with anti-CTCF (2 μg, 07-729, Millipore) or anti-HNF4α (2 μg, ab181604, Abcam) antibodies in dilution buffer (16.7 mM Tris-HCl pH 8.1, 167 mM NaCl, 0.01% SDS, 1.1% Triton-X 100, protease inhibitor) at 4°C overnight. Protein-A agarose beads were also washed with cold dilution buffer three times and incubated with blocking buffer (10 mg/ml BSA, ChIP dilution buffer, protease inhibitor) at 4°C overnight. Sheared DNA incubated with antibody and blocked protein-A agarose were incubated at 4°C for one hour the next day and washed at room temperature with buffers TSEI (20 mM Tris-HCl pH 8.1, 150 mM NaCl, 2 mM EDTA, 0.1% SDS, 1% Triton X-100), TSE II (20 mM Tris-HCl pH 8.1, 500 mM NaCl, 2 mM EDTA, 0.1% SDS, 1% Triton X-100), ChIP buffer III (10 mM Tris-HCl pH 8.1, 0.25M LiCl, 1 mM EDTA, 1% NP-40, 1% deoxycholate), and TE (10 mM Tris-HCl pH 8.1, 1 mM EDTA). Chromatin was eluted with elution buffer (1% SDS, 0.1 M NaHCO_3_) twice and incubated with 0.2 M NaCl at 65 °C overnight to reverse the cross links. Digestion was carried out with 10 mg/mL proteinase K in 40 mM Tris-HCl pH 7.5 and 10 mM EDTA to purify CTCF-or HNF4α-bound and input DNA. ChIP-seq libraries were prepared with the NEBNext Ultra II DNA Library Prep Kit for Illumina (E7645S, New England BioLabs) and Agencourt AMPure XP beads were used for size selection to generate the final libraries. Library quality was assessed with an Agilent High Sensitivity DNA Bioanalyzer (5067-4626, Agilent Technologies), and quantity measured with KAPA Library Quantification Kits (KK4835, KAPA Biosystems).

### ChIP-seq data analysis

ChIP-seq libraries were sequenced on an Illumina HiSeq 4000 (Illumina) with 100 single-end reads and aligned to the mm10 genome with STAR^89^. Bam files from various technical replicates of the same biological sample were merged with Samtools^82^. Peak calling was performed with Homer^90^ and differential occupancy analysis was carried out with the R package DiffBind^86^. Read counts for each peak were quantified with dba.count (score=DBA_SCORE_TMM_MINUS_FULL, bUseSummarizeOverlaps=TRUE) and differential analysis were identified with dba.analyze (method=DBA_EDGER, bSubControl=T, bTagwise=F) in conjunction with edgeR^87^.

### CTCF differential expression insulator analysis

Increased CTCF occupancy during liver repopulation could prevent distal regulatory regions to activate only one of the flanking promoters surrounding a CTCF binding site, and therefore leading to a larger difference in gene expression levels. We define this ‘differential expression insulator’ function, in which a gene pair is either highly or lowly expressed without the presence of CTCF, but only one flanking gene exhibits a decrease in gene expression after binding of CTCF. An insulator strength score was calculated for all significantly gained (fold change ≥1.5, FDR ≤0.05) CTCF peaks in the repopulating liver as previously described^51^. Briefly, CTCF sites with divergent flanking promoters within 50 kb were identified and the corresponding gene expression levels from quiescent and 4-week repopulating hepatocytes were extracted from published TRAP-seq^15^.

Low-expressors, in which RPKM-normalized read counts are 0 across all samples, were filtered followed by calculation of a rank percentile based on RPKM for each gene. Let *x*_*Q*_ and *y*_*Q*_ be the expression percentile in the quiescent hepatocytes; *x*_*R*_ and *y*_*R*_ be the expression percentile in the 4-week repopulating hepatocytes. The insulator strength score is calculated by taking the maximum value of *x*_*Q*_ × *y*_*Q*_ × *x*_*R*_ × *1−yR* and *xQ*×*yQ*×*1−xR*×*yR*. A differential expression insulator function will have one of the following effects: (1) Increased *x*_*R*_ and decreased *y*_*R*_: in this case, *x*_*Q*_ × *y*_*Q*_ × *x*_*R*_ × (1-*y*_*R*_) will be the largest. (2) Decreased *x*_*R*_ and increased *y*_*R*_: in this case, *x*_*Q*_ × *y*_*Q*_ × (1 -*x*_*R*_) × *y*_*R*_ will be the largest. Gained CTCF sites with the top 25% insulator strength scores were categorized as strong insulators. Random gene pairs not flanked by CTCF within 50 kb were used as controls and a differential expression insulator score for each gene pair was calculated as described above. The number of significant (FDR≤0.05) and non-significant (FDR>0.05) differential expression of the flanking genes were identified for all strong insulators from increased CTCF binding and random genomic regions. Finally, we used a Fisher’s exact test to examine the likelihood of gained CTCF sites to contain more significantly changed genes when compared to that of control regions.

### Nucleosome location analysis with ATAC-seq

MAC2 (callpeak --keep-dup all, -B --SPMR, -q 0.05, --broad)^84^ was used to identify broad peaks from all aligned bam files including nucleosome-free reads and nucleosome-containing reads from ATAC-seq. Broad peaks were then processed with BEDtools^92^ to extend the peaks (bedtools slop -b 200), sorted by genomic positions (sort -k1,1 -k2,2n), and overlapping reads were merged (bedtools merge). Nucleosome position was identified with NucleoATAC^93^ from the aligned bam and broad peak files. The closest nucleosomes with respect to TSS were identified, and those within 350 bp upstream and 250 bp downstream of the TSS were identified as the −1 and +1 nucleosomes, respectively.

### Nucleosome positioning analysis

The distance of +1 to −1 nucleosomes was calculated for each transcript. We used the Kolmogorov-Smirnov test to compare the +1 and −1 nucleosome distribution differences between quiescent and repopulating hepatocytes, respectively. To analyze the association between gene activity and nucleosome positioning, transcriptomic changes in repopulating hepatocytes^15^ were first stratified into three categories: top 500 upregulated (fold change ≥1.5, FDR ≤0.05), top 500 downregulated (fold change ≥1.5, FDR ≤0.05), and unchanged (absolute fold change <1.5 or FDR >0.05) genes. The distances between the +1 to −1 nucleosomes were calculated for each gene and differential positioning was carried out by comparing the distance in quiescent to regenerating hepatocytes in the upregulated, downregulated, and unchanged gene expression groups, respectively, with a permutation test (n=10,000).

### Statistical analysis

EdgeR^87^ was used for all high-throughput sequencing data analysis. For the integrative TRAP-seq and ATAC-seq analysis, a hypergeometric test was used for identifying the significance of overlapping gene sets, and a one-sample t test was used to compare the difference between normalized gene expression fold change in DA promoter and enhancer peaks, respectively. A Kolmogorov-Smirnov test was performed for global distribution change of +1 and −1 nucleosome positioning and a permutation test (n=10,000) was carried out to test the change in +1 to −1 nucleosome distance of genes with differential expression.

### Study approval

The animal experiments carried out in this study were reviewed and approved by the IACUC of the Penn Office of Animal Welfare at the University of Pennsylvania.

### Access to data

All authors had access to the study data and had reviewed and approved the final manuscript.

## Disclosures

The authors have nothing to disclose

### Transcript and Chromatin Profiling

All sequencing data in this study were deposited to the Gene Expression Omnibus database and can be downloaded with the accession number GSE109466

### Author Contributions

AWW and YJW acquired the ATAC-seq data. AWW performed data analysis, interpretation, statistical analysis, and wrote the manuscript. AMZ assisted with data interpretation and statistical analysis. ARM performed all ChIP-seq experiments. KJW conducted all mouse injection experiments. KHK supervised the study. All authors edited the manuscript.

